# Occurrence, incidence, and severity of Alternaria leaf spot of cotton (*Gossypium hirsutum L.*) In major cotton-growing districts of the Lake Zone, Tanzania

**DOI:** 10.64898/2026.09.24.754005

**Authors:** Salvatory Mwesige Eustace, Beatrice V. Mwaipopo, Richard Raphael Madege

## Abstract

Alternaria leaf spot (ALS), caused by fungi in the genus Alternaria (family *Pleosporaceae*), is a significant fungal disease affecting cotton production in the Lake Zone of Tanzania, the country’s main cotton-producing region, where it reduces photosynthetic capacity, boll retention, and fiber quality. Despite its increasing prevalence, systematic baseline epidemiological data remain scarce. This study aimed to determine the occurrence, incidence, and severity of Alternaria leaf spot across the major cotton-growing districts of the Lake Zone, Tanzania. Field surveys were conducted during the 2024-2025 growing season in Bariadi, Bunda, Chato, and Magu districts; 20 representative cotton fields were systematically assessed using a diagonal sampling approach, and disease incidence, prevalence, and severity were recorded for 1,000 cotton plants. Fungal isolates recovered from symptomatic leaves were examined for morphological identification. Field prevalence was 100%, and overall mean incidence was 64.4%, with significant inter-district variation: Bariadi (77.6%), Bunda (68.8%), Chato (60.0%), and Magu (52.4%). Mean disease severity index (DSI) ranged from 44.0% (Magu) to 70.1% (Bariadi), indicating moderate-to-severe infection. Incidence and DSI were strongly correlated when uninfected plants were included (r = 0.942, P < 0.001), but this relationship weakened and became non-significant when restricted to infected plants only (r = 0.396, P = 0.084), suggesting that higher incidence reflected more plants becoming infected rather than greater severity among already-infected plants. Morphological characterization identified all 40 isolates as belonging to the genus Alternaria. These findings provide the first systematic epidemiological baseline for Alternaria leaf spot across the major cotton-growing districts of Tanzania’s Lake Zone. Disease incidence and severity varied significantly across districts, underscoring the need for tailored disease management strategies; this study recommends ongoing surveillance and integrated disease management to safeguard cotton productivity among smallholder farmers in the region.

## 1.0 INTRODUCTION

Cotton (*Gossypium hirsutum L*.), commonly known as “white gold,” is one of the world’s most important fibre crops and plays a central role in the global textile industry. It is cultivated in more than 80 countries and provides income and employment to millions of people involved in production, processing, and marketing (Kaliyaperumal & Rajasekaran, 2023). Cotton contributes significantly to the crop value chain by generating employment, supporting textile manufacturing, generating export revenue, and supplying edible oil and animal feed (FAO, 2022). Major cotton-producing countries, including China, India, the United States, Brazil, and Pakistan, together account for more than 70% of global production (FAO, 2022). In Africa, cotton serves as a strategic cash crop, particularly in West, Central, and Eastern Africa, where it supports rural livelihoods and national economies (Jans *et al*., 2022; Soumaré *et al*., 2021). Among Eastern African nations, Tanzania stands out as a key cotton producer, where the crop plays a similarly vital role in the national economy and smallholder farming systems.

In Tanzania, cotton is the second most important export crop after coffee, contributing 3-5% of national GDP and supporting approximately 600,000 smallholder farming households annually (URT, 2024). During the 2023-2024 season, total production reached 149,361 tonnes, and lint exports generated between USD 80 million and USD 380 million in foreign exchange earnings (NBS, 2024). Cotton is predominantly produced in the Lake Zone, comprising the Simiyu, Mara, Mwanza, and Geita regions, which together form the Western Cotton-Growing Areas (WCGA) (Ministry of Agriculture, 2024). These regions are characterized by favourable agroecological conditions, including moderate rainfall, suitable temperatures, and relatively fertile soils, which support sustainable cotton production (Omar, 2023).

Despite these favourable conditions, however, cotton production in Tanzania has been declining due to various constraints, including drought, insect pests, diseases, weeds, declining soil fertility, and inadequate input distribution systems (Faustine *et al*., 2016; Mung’ong’o, 2024). According to the Ministry of Agriculture (2024), cotton productivity in Tanzania declined to 1.34 tonnes per hectare, significantly below the potential yield of 3 tonnes per hectare, signaling a substantial productivity crisis that undermines both farmers’ incomes and national economic growth. Among the biotic constraints, fungal diseases such as Alternaria leaf spot, bacterial blight, and Fusarium wilt have been reported to cause significant losses (Hillocks, 1991; Lawrence *et al*., 2014; Wang, 2024).

Among these three diseases, Alternaria leaf spot, caused mainly by *Alternaria macrospora Zimm*. and *A*. *alternata (Fr.) Keissl*., has emerged as a major foliar disease threatening cotton production. The disease affects all stages of cotton development, primarily targeting late-season or aging tissues. Characteristic symptoms include small, circular to irregular brown necrotic lesions with concentric rings, chlorotic halos, and premature leaf senescence. Over time, lesions expand, becoming dry, gray, and shot-holed, thereby reducing photosynthetic capacity and cotton yield (Lin *et al*., 2024). High disease incidence and severity are favoured by elevated humidity, frequent rainfall, and prolonged leaf wetness (Bhat *et al*., 2013; Kumar *et al*., 2018). Globally, the disease has been documented as a significant constraint in major cotton-producing countries, including Mexico, China, and India (Li *et al*., 2023; Zhu et al., 2019; Sampathkumar *et al*., 2023). In Africa, Alternaria leaf spot has been reported in Zimbabwe, Sudan, and Egypt (Hillocks, 1991; Mohamed et al., 2019; Olmez *et al*., 2023), causing yield losses exceeding 40% and accompanied by reduced fiber quality and boll retention.

Similarly, in Tanzania, Alternaria leaf spot has increasingly been observed in cotton fields within the Lake Zone, where cotton is predominantly grown under rainfed smallholder systems, with documented yield losses of 30-40% under favourable environmental conditions (Mung’ong’o, 2024; URT, 2024). Despite these mounting losses and the visible spread of the disease across the WCGA, critical information gaps persist regarding the incidence, severity, and inter-district variability of the disease across major cotton-growing areas. The Lake Zone, while recognized as the epicentre of Tanzanian cotton production and increasingly affected by Alternaria leaf spot, has received minimal systematic disease surveillance. This represents a critical knowledge gap because: (i) smallholder farmers lack region-specific management guidance, (ii) extension services cannot prioritize resources without understanding disease distribution, and (iii) breeding programs require baseline incidence data for variety selection. Without baseline epidemiological data on disease occurrence and its relationship with environmental factors, farmers and extension services lack evidence-based guidance for disease monitoring and control.

This study therefore aimed to: (i) determine the occurrence and prevalence of Alternaria leaf spot across the major cotton-growing districts of the Lake Zone; (ii) quantify disease incidence and severity at the district and village levels; (iii) conduct preliminary morphological identification of Alternaria isolates recovered from symptomatic cotton plants

## 2.0 MATERIALS AND METHODS

### 2.1 Study Area

The study was conducted in four major cotton-growing districts of the Lake Zone of Tanzania: Bariadi District (Simiyu Region), Bunda District (Mara Region), Chato District (Geita Region), and Magu District (Mwanza Region) (Fig. 1). These districts represent distinct cotton production agroecologies and differ in rainfall patterns, temperature regimes, and agronomic practices that may influence the epidemiology of Alternaria leaf spot. Bariadi District lies between approximately 2.80 °- 3.50 ° S and 33.80 °- 35.00 ° E; Bunda District lies between 1.80 °- 2.50 ° S and 33.80 °- 34.50 ° E; Chato District lies between 2.50°–3.50° S and 31.00°– 32.00° E; and Magu District lies between 2.00°–3.00° S and 33.00 °- 34.00 ° E. All study areas are characterized by tropical savannah climatic conditions with distinct wet and dry seasons, annual rainfall ranging from 700 to 1,200 mm, and temperatures between 20°C and 32°C.

**Figure 1:**
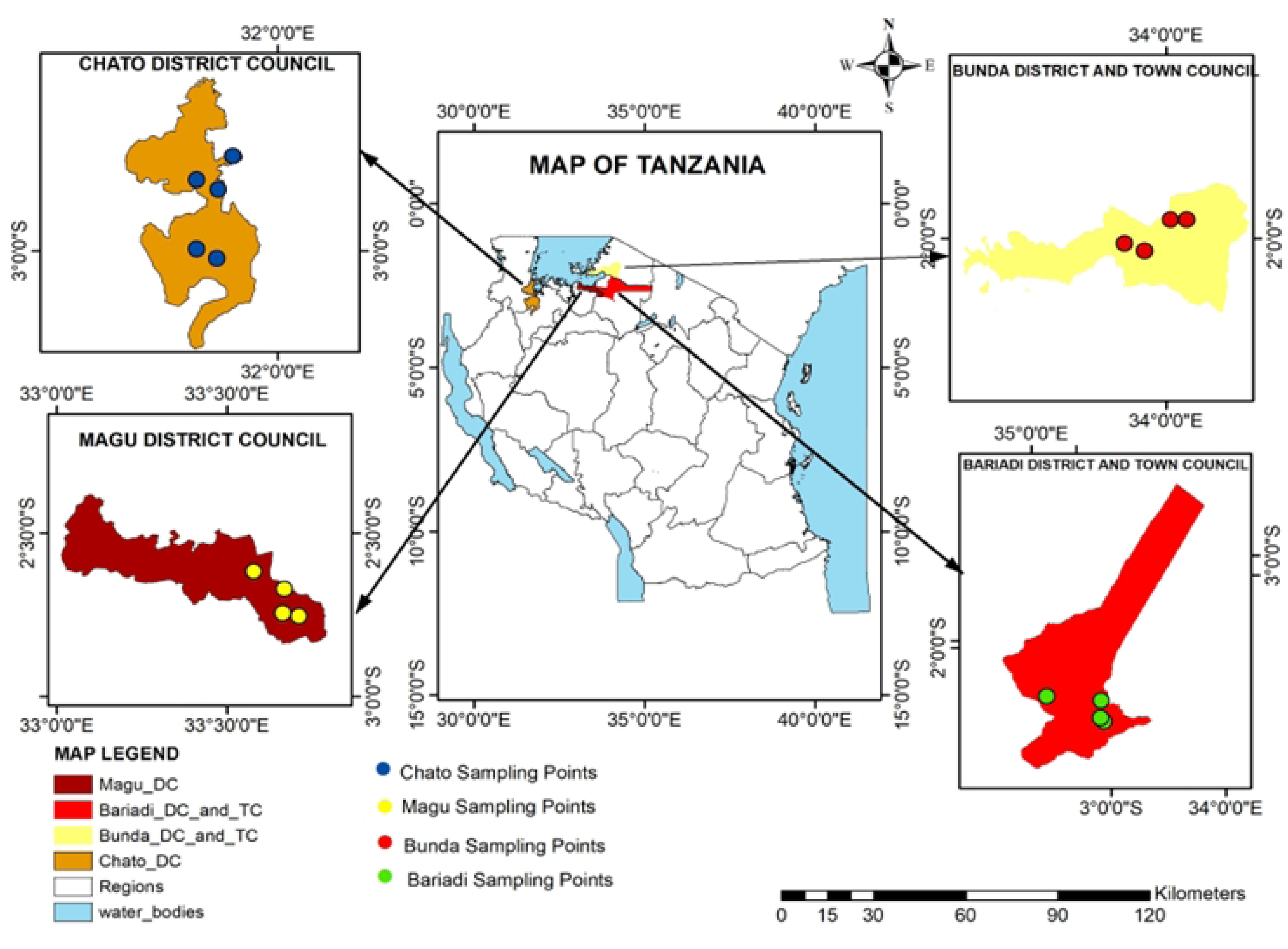
Map of Tanzania showing the four cotton-growing districts surveyed in this study: Bariadi (Simiyu Region), Bunda (Mara Region), Chato (Geita Region), and Magu (Mwanza Region).

### 2.2 Study design and sampling

A cross-sectional observational design was used, employing a purposive multistage sampling technique as described by Kothari (2004). Districts were selected based on cotton production potential and reported disease incidence, with logistical support from District Agricultural, Livestock, and Fishery Officers (DALFOs). Within each of the four selected districts, five wards and their respective villages were purposively selected. Field surveys were conducted from March to May 2025, corresponding to the late vegetative, flowering, and boll development stages of cotton growth, when Alternaria leaf spot symptoms are most pronounced. A total of 20 cotton fields (five per district), each at least one hectare, were surveyed, with one representative field selected per village. Surveyed fields were spaced at 5- 10 km intervals to ensure geographic coverage and minimize sampling overlap.

An equal-allocation design was adopted across the four major cotton-producing districts of the Lake Zone, rather than proportional allocation, given the logistical constraints inherent to a first systematic baseline survey in which no prior epidemiological data existed to inform weighting. Field-level replication (five fields per district, n = 20 total) was purposefully paired with intensive within-field sampling (50 plants per field, 1,000 plants overall) so that each field-level incidence and severity value represents a precise estimate rather than a single observation, partially offsetting the modest number of field replicates.

The survey prioritized broad geographic coverage to capture agroecological variation and establish baseline data for future epidemiological studies.

Within each field, 50 plants were assessed: 10 plants were randomly selected from each of the four corners (southwest, southeast, northwest, and northeast) and from the middle of the field, using a diagonal sampling method to enhance spatial representativeness within the field and minimize sampling bias (Madden *et al*., 2007).

### 2.3 Disease identification and assessment

Disease identification and assessment followed standard phytopathological procedures for field-based disease surveys. Typical symptoms of Alternaria leaf spot, including circular to irregular necrotic lesions with concentric ring patterns, chlorotic halos, and premature leaf senescence, served as the basis for field diagnosis, guided by established symptom descriptions (Bhat *et al*., 2013; Lin *et al*., 2024). High-quality images of infected plant parts were captured under natural light with a digital camera for documentation. Only leaves showing clear, typical symptoms were included in laboratory analysis; leaves with mixed or ambiguous symptoms were excluded to minimize diagnostic error. Asymptomatic leaves collected from surveyed fields served as controls.

Disease incidence was determined by counting the number of infected plants relative to the total number of plants assessed in each field. Incidence was expressed as a percentage using the standard formula (Teng & James, 2002).

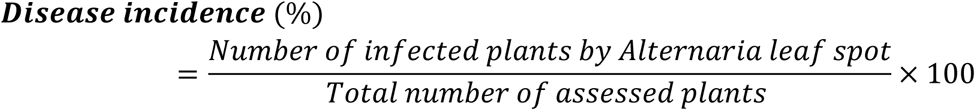

Disease prevalence was assessed at the village and district levels by recording the number of fields with Alternaria leaf spot. Prevalence was expressed as the percentage of infected fields relative to the total number of surveyed fields (Sanogo & Carpenter, 2006).

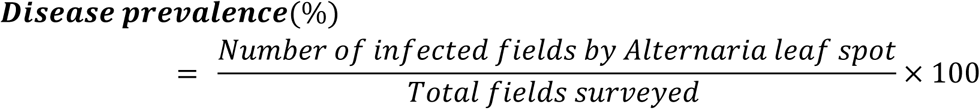

Cotton Alternaria leaf spot disease severity was assessed in the surveyed fields using a 0–5 rating scale as described by Bhat et al. (2013). Each plant was visually examined, and the percentage of leaf area infected was recorded on the scale (Table 1). To standardize severity across fields, the Disease Severity Index (DSI) was calculated using the formula (Park et al., 2008):

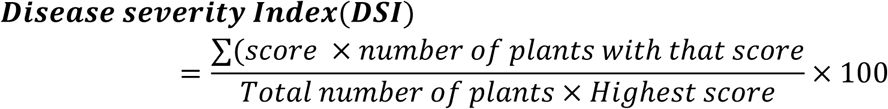

**Table 1.** Disease severity rating scale for Alternaria leaf spot of cotton.

| Score | Description of symptoms | % Infected leaf area |
| --- | --- | --- |
| 0 | No disease symptoms on the leaves | 0% |
| 1 | Very mild infection | 0.1 – 10.0% |
| 2 | Mild infection | 10.1 – 25.0% |
| 3 | Moderate infection | 25.1 – 50.0% |
| 4 | Severe infection | 50.1 – 75.0% |
| 5 | Very severe infection | >75.0% |

This provided a percentage value representing the overall severity of infection in each field.

### 2.4 Laboratory isolation and morphological identification

After sampling, samples were transported in sealed polythene bags on ice and processed within 24 hours to minimize tissue deterioration and contamination (Ahmad *et al*., 2024). Leaf sections (5 × 5 mm) were excised from lesion margins with a sterile scalpel, ensuring each section included both symptomatic and adjacent asymptomatic tissue. Sections were sequentially surface sterilized with 70% ethanol (30 s), 1% sodium hypochlorite (1 min), and sterile distilled water (3 rinses), following established fungal isolation protocols (Dhingra & Sinclair, 1995). Sterilized tissues were plated on potato dextrose agar (PDA) amended with streptomycin sulfate (100 mg L⁻¹) to suppress bacterial contamination and incubated at 25 ± 2°C under a 12-hour photoperiod for 7–10 days to promote fungal growth (Zhu *et al*., 2019). Primary isolation plates typically yielded mixed fungal growth; colonies exhibiting Alternaria-like morphology (dark olive-gray, velvety texture, irregular margins) were selected, and emerging hyphal tips were subculture onto fresh PDA to obtain pure cultures. These were further purified to obtain genetically uniform isolates for morphological characterization and were subsequently preserved in sterile distilled water at 4°C for future use (Ahmad *et al*., 2024).

Morphological identification was performed on pure cultures obtained from hyphal tips grown on PDA for 7 days at 25°C, a temperature considered optimal for Alternaria sporulation and colony development (Anil et al., 2017; Sampathkumar *et al*., 2024). Colony features, including colour, texture, growth pattern, and reverse pigmentation, were recorded using standard mycological procedures (Simmons, 2007). Microscopic observations of conidia and conidiophores were made from lactophenol cotton blue mounts under a compound microscope (Olympus CX23, Olympus Corporation, Tokyo, Japan). Measurements of at least 30 conidia per isolate were recorded, including conidial length, width, number of transverse and longitudinal septa, and beak length (Nicolai *et al*., 2023). Morphological identification was used to confirm that isolates belonged to the genus *Alternaria*.

**Figure 2:**
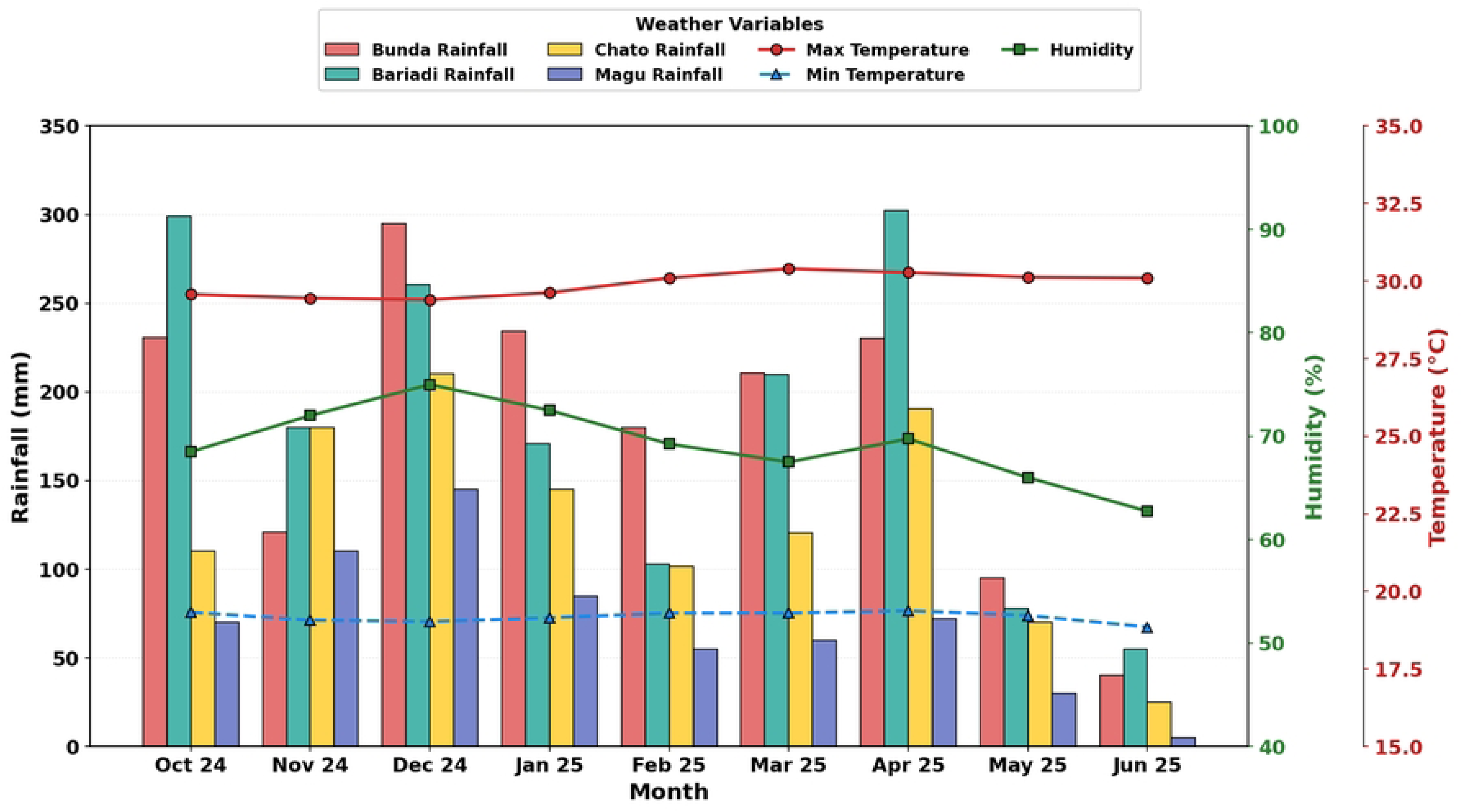
Seasonal climatic data for the four surveyed districts during the 2024-2025 cotton-growing season (Source: Tanzania Meteorological Authority).

### 2.5 Climatic data collection

Seasonal rainfall, mean maximum and minimum temperatures, and mean relative humidity (RH) for each surveyed district during the 2024-2025 growing season were obtained from the Tanzania Meteorological Authority (TMA). These district-level climatic data (Figure 3) were used to characterize the prevailing environmental conditions during the survey period.

**Figure 3:**
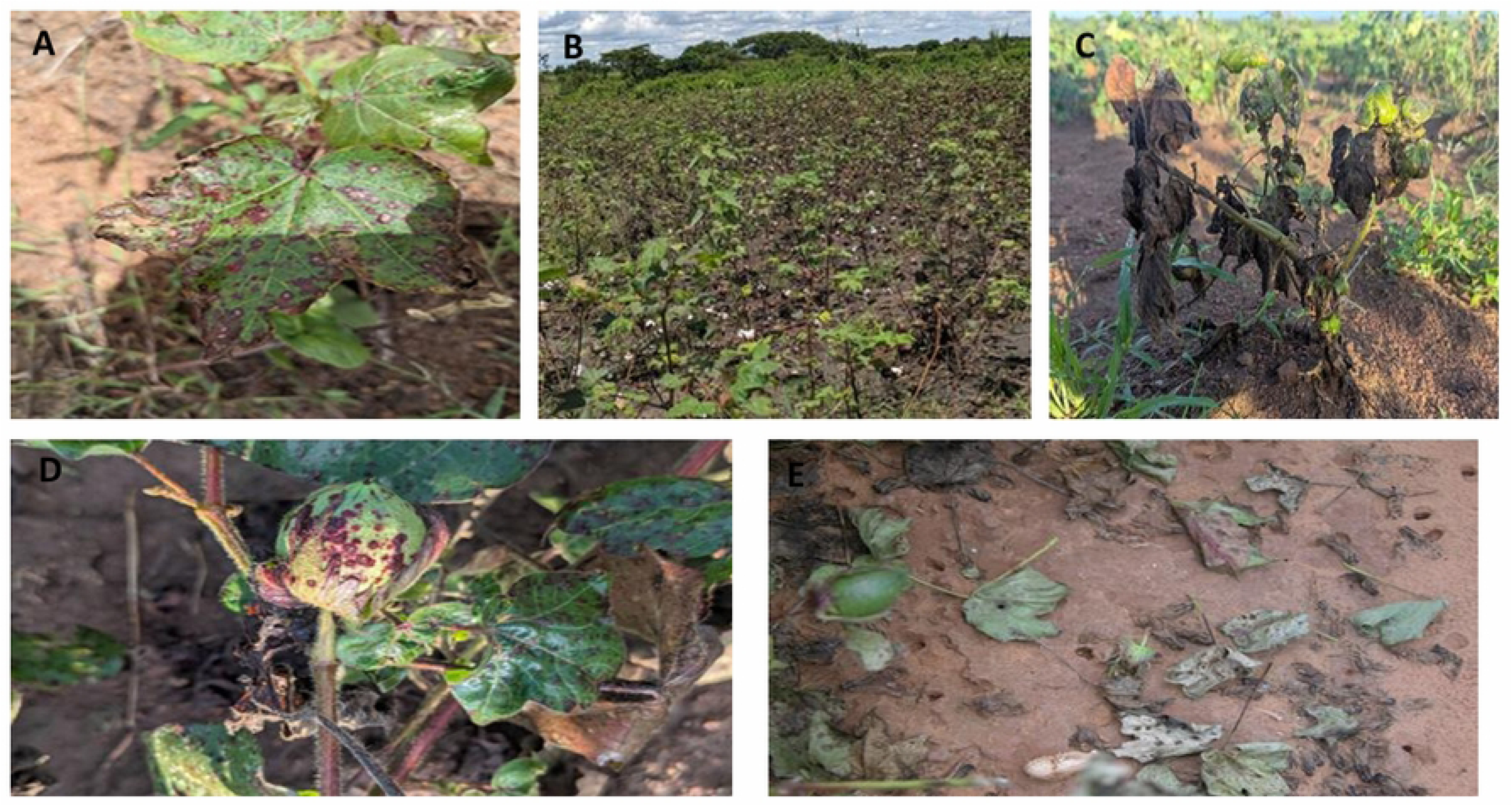
Symptoms of Alternaria leaf spot of cotton observed during field surveys in the Lake Zone of Tanzania: (A) Circular necrotic lesions with concentric ring patterns on a cotton leaf; (B) Cotton field exhibiting widespread Alternaria leaf spot incidence; (C) Cotton plant showing severe leaf defoliation at advanced disease stage; (D) Cotton bowl with necrotic lesions caused by Alternaria spp.(E) Defoliated cotton leaves associated with severe Alternaria infection.

### 2.6 Data Analysis

Descriptive statistics (means, minimum and maximum values, standard deviations, and coefficients of variation) were computed to summarize the distribution and intensity of Alternaria leaf spot across surveyed districts, wards, and fields. Before analysis, data were assessed for normality using the Shapiro–Wilk test and for homogeneity of variance using Levene’s test. Disease incidence data from districts with unequal variances were arcsine-transformed to stabilize variance and improve normality. One-way analysis of variance (ANOVA) was performed to assess significant differences in disease incidence, mean severity score (MSS), and DSI among districts, followed by Tukey’s Honest Significant Difference (HSD) test for pairwise mean comparisons at P < 0.05. Untransformed means and standard deviations are presented in tables and text for biological interpretation. Ninety-five percent confidence intervals (95% CI) were calculated for mean incidence, mean severity score, and DSI at the district level to indicate the precision of each estimate; for field-level prevalence, exact (Clopper-Pearson) binomial 95% CIs were calculated. To quantify the relationship between disease incidence and DSI across all 20 surveyed fields, Pearson product-moment correlation and simple linear regression (DSI as the response variable and incidence as the predictor) were performed. Spearman’s rank correlation was computed as a non-parametric verification. Village-level data (one field per village) were presented descriptively only, as the single-observation-per-village design precluded inferential comparison among villages within districts. Statistical analyses were performed using Python 3.11 (SciPy v1.11; stats models) for descriptive statistics and correlation analyses, and R version 4.4.2 (R Core Team, 2024) for ANOVA and regression analyses. The significance level was set at P < 0.05 for all tests.

## 3.0 RESULTS

### 3.1 Disease prevalence

Alternaria leaf spot symptoms, including circular to irregular necrotic lesions with distinct concentric ring patterns and, in severe cases, premature leaf senescence, were observed in all surveyed cotton fields (Figure 3). Field-level prevalence, defined as the percentage of surveyed fields with symptomatic plants, Alternaria leaf spot was detected in all 20 surveyed fields across the four districts, corresponding to 100% field-level prevalence.

### 3.2 Disease incidence of Alternaria leaf spot

Disease incidence varied significantly among districts at P < 0.001. The overall mean disease incidence was 64.4% (95% CI: 59.5-69.3%). The highest incidence was recorded in Bariadi District (77.6 ± 4.8%; 95% CI: 71.2-83.2%), followed by Bunda (68.8 ± 5.7%; 95% CI: 60.5-74.7%), Chato (60.0 ± 5.7%; 95% CI: 52.9-67.1%), and Magu (52.4 ± 4.8%; 95% CI: 46.5-58.3%). Tukey’s HSD test showed that Bariadi differed significantly from Chato (P = 0.0004) and Magu (P < 0.0001), while Bunda differed significantly from Magu (P = 0.0009), whereas Bariadi and Bunda did not differ significantly from each other (P = 0.079), nor did Bunda and Chato (P = 0.079), nor did Chato and Magu (P = 0.149; Figure 4).

**Figure 4:**
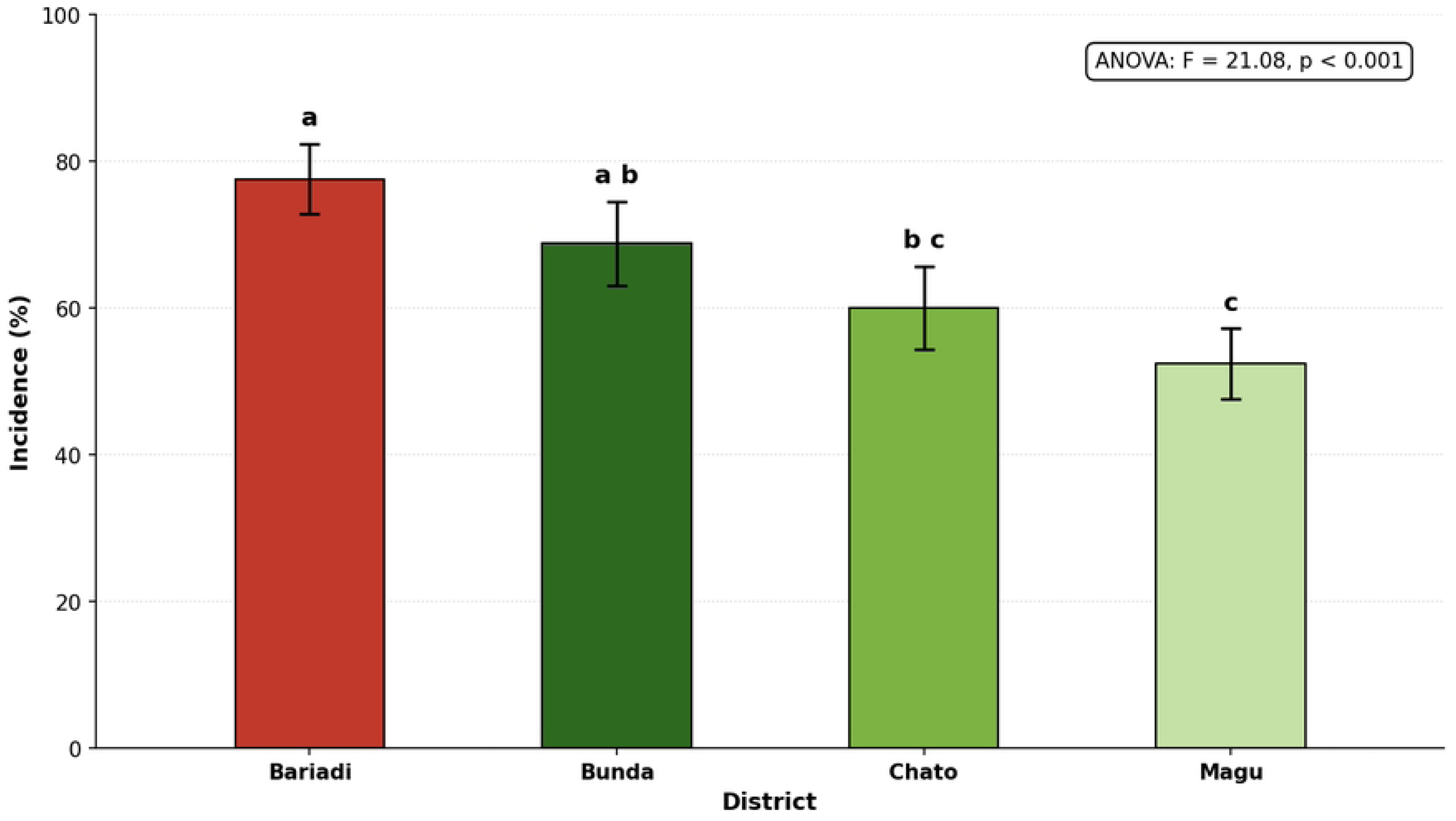
Disease incidence (%) of Alternaria leaf spot across the four surveyed districts of the Lake Zone of Tanzania. Error bars represent ±SE (n=5 fields per district). Different letters indicate significant differences among districts (Tukey HSD, α = 0.05). ANOVA: F = 21.08, p < 0.001.

### 3.3 Disease severity of *Alternaria* leaf spot

Disease severity varied significantly across districts (P < 0.05). The highest mean severity was recorded in Bariadi District (MSS = 3.50, 95% CI: 3.21–3.79; DSI = 70.0%, 95% CI: 64.1-75.9%), followed by Bunda (MSS = 3.12, 95% CI: 2.96-3.28; DSI = 62.4%, 95% CI: 59.2-65.6%), Chato (MSS = 2.50, 95% CI: 2.26-2.74; DSI = 50.0%, 95% CI: 45.3-54.7%), and Magu (MSS = 2.21, 95% CI: 2.09-2.33; DSI = 44.2%, 95% CI: 41.8-46.6%). The overall mean DSI was 56.7% (95% CI: 51.6-61.7%), indicating a moderate-to-severe infection across the surveyed production areas. A clear gradient in disease intensity was observed, with Bariadi and Bunda experiencing significantly greater disease pressure than Chato and Magu.

### 3.4 Village-level variation in disease incidence and severity

Considerable within-district variability in disease incidence and DSI was observed at the village level across all four districts (Table 4). Village-level data were presented descriptively because the sampling design allocated one field per village (n = 1), precluding inferential comparisons among villages within a district. Within-district variability statistics are shown in Figure 4 for incidence and in Table 3 and Figure 5 for severity.

**Figure 5:**
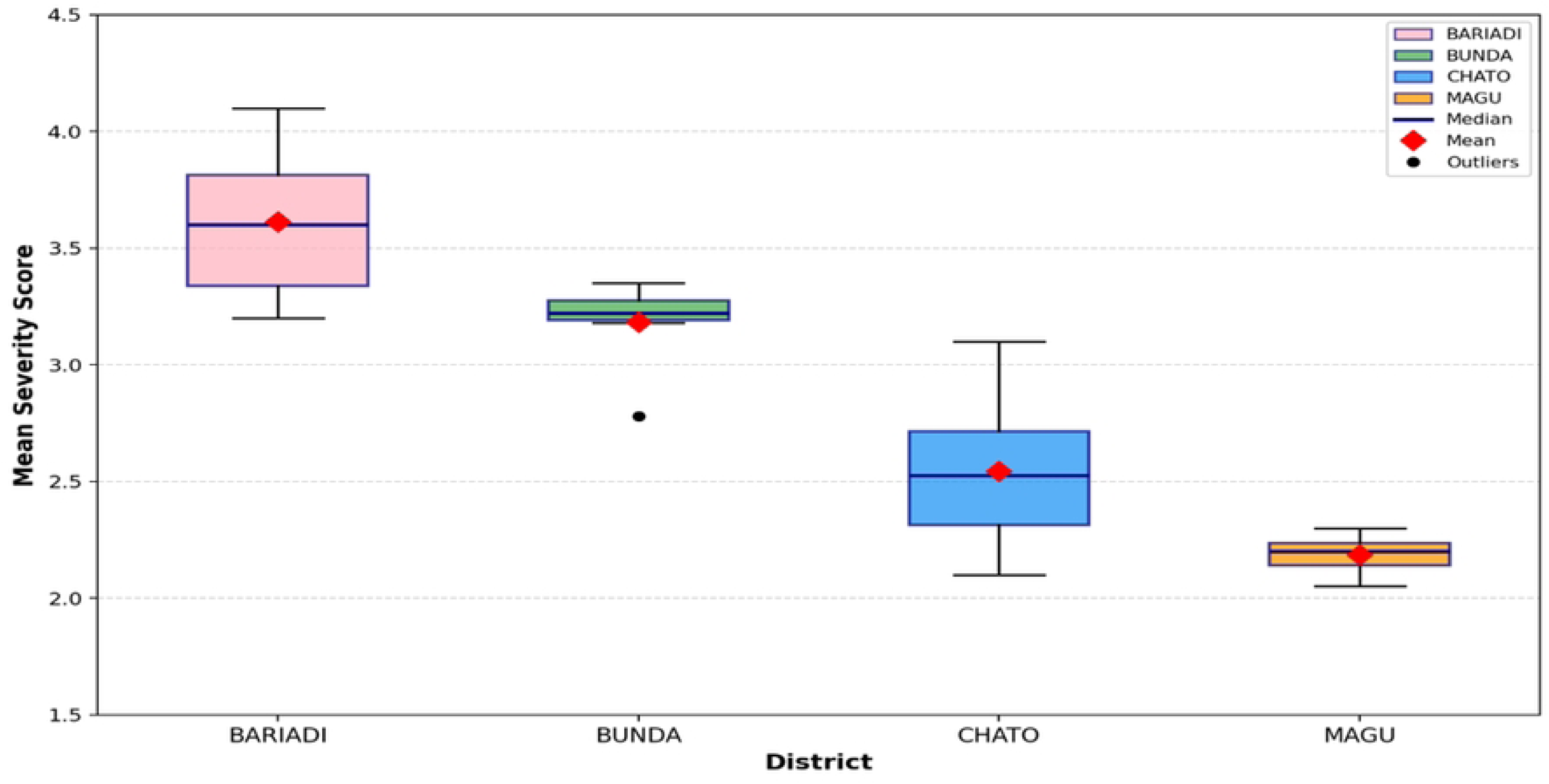
Box plot of the mean severity score of Alternaria leaf spot across surveyed districts. Boxes represent interquartile range; horizontal line = median; whiskers = full data range; dots = Outlier

**Table 3.** Mean severity score (MSS), Disease Severity Index (DSI), and coefficient of variation (CV) of Alternaria leaf spot across surveyed districts in the Lake Zone, Tanzania.

| District | Field surveyed | MSS | SE | DSI (%) | SE | CV (%) |
| --- | --- | --- | --- | --- | --- | --- |
| **Bariadi** | 5 | 3.50<sup>a</sup> | 0.16 | 70.1<sup>a</sup> | 3.35 | 10.7 |
| **Bunda** | 5 | 3.13<sup>b</sup> | 0.09 | 62.6<sup>b</sup> | 1.74 | 6.2 |
| **Chato** | 5 | 2.50<sup>c</sup> | 0.17 | 50.1<sup>c</sup> | 3.53 | 15.8 |
| **Magu** | 5 | 2.20<sup>c</sup> | 0.05 | 44.0<sup>c</sup> | 1.12 | 5.7 |
| **Mean** |  | **2.83** |  | **56.7** |  |  |

Means followed by different superscript letters within a column differ significantly at $P < 0.05$ according to Tukey's HSD test. MSS = Mean Severity Score; DSI = Disease Severity Index; CV = Coefficient of variation; SE = Standard error. Pairwise Tukey's HSD P-values (identical for MSS and DSI): Bariadi vs Bunda, $P = 0.223$ ; Bariadi vs Chato, $P = 0.0003$ ; Bariadi vs Magu, $P < 0.0001$ ; Bunda vs Chato, $P = 0.017$ ; Bunda vs Magu, $P = 0.0006$ ; Chato vs Magu, $P = 0.381$ .

**Table 4.** Village-level disease incidence and severity of Alternaria leaf spot across surveyed districts in Tanzania’s Lake Zone.

| District | Village | Incidence (%) | MSS | DSI (%) |
| --- | --- | --- | --- | --- |
| **Bariadi** | Mbalanga | 84.0 | 3.80 | 76.0 |
|  | Missi | 80.0 | 3.65 | 73.0 |
|  | Zina | 76.0 | 3.50 | 70.0 |
|  | Salmu | 74.0 | 3.35 | 67.0 |
|  | Nyakabindi | 72.0 | 3.20 | 64.0 |
| **Bunda** | Makongeni | 74.0 | 3.30 | 66.0 |
|  | Bushigwamhala | 72.0 | 3.20 | 64.0 |
|  | Kunzungu | 68.0 | 3.10 | 62.0 |
|  | Ichamu | 64.0 | 3.00 | 60.0 |
|  | Tairo | 60.0 | 3.00 | 60.0 |
| **Chato** | Bwanga | 68.0 | 2.75 | 55.0 |
|  | Bukiriguru | 64.0 | 2.60 | 52.0 |
|  | Busaka | 60.0 | 2.50 | 50.0 |
|  | Bwera | 56.0 | 2.40 | 48.0 |
|  | Rwantaba | 54.0 | 2.25 | 45.0 |
| **Magu** | Mwamibanga | 58.0 | 2.35 | 47.0 |
|  | Nyanshoshi | 56.0 | 2.25 | 45.0 |
|  | Nyang'hanga | 52.0 | 2.20 | 44.0 |
|  | Ng'haya | 50.0 | 2.15 | 43.0 |
|  | Misungwi | 46.0 | 2.10 | 42.0 |

Note: Village-level values are recorded per individual field. MSS = Mean Severity Score; DSI = Disease Severity Index.

### 3.5 Relationship between disease incidence and severity

Disease incidence and mean severity across all assessed plants showed a strong positive correlation (r = 0.942, P < 0.001), as both parameters were derived from the same disease severity ratings. To determine whether this relationship represented a biological association, mean severity was recalculated using only infected plants, excluding healthy plants (class 0). The correlation became weaker and non-significant (r = 0.396, P = 0.084; Figure 6), indicating that the proportion of infected plants was not necessarily associated with disease intensity within the infected population.

**Figure 6:**
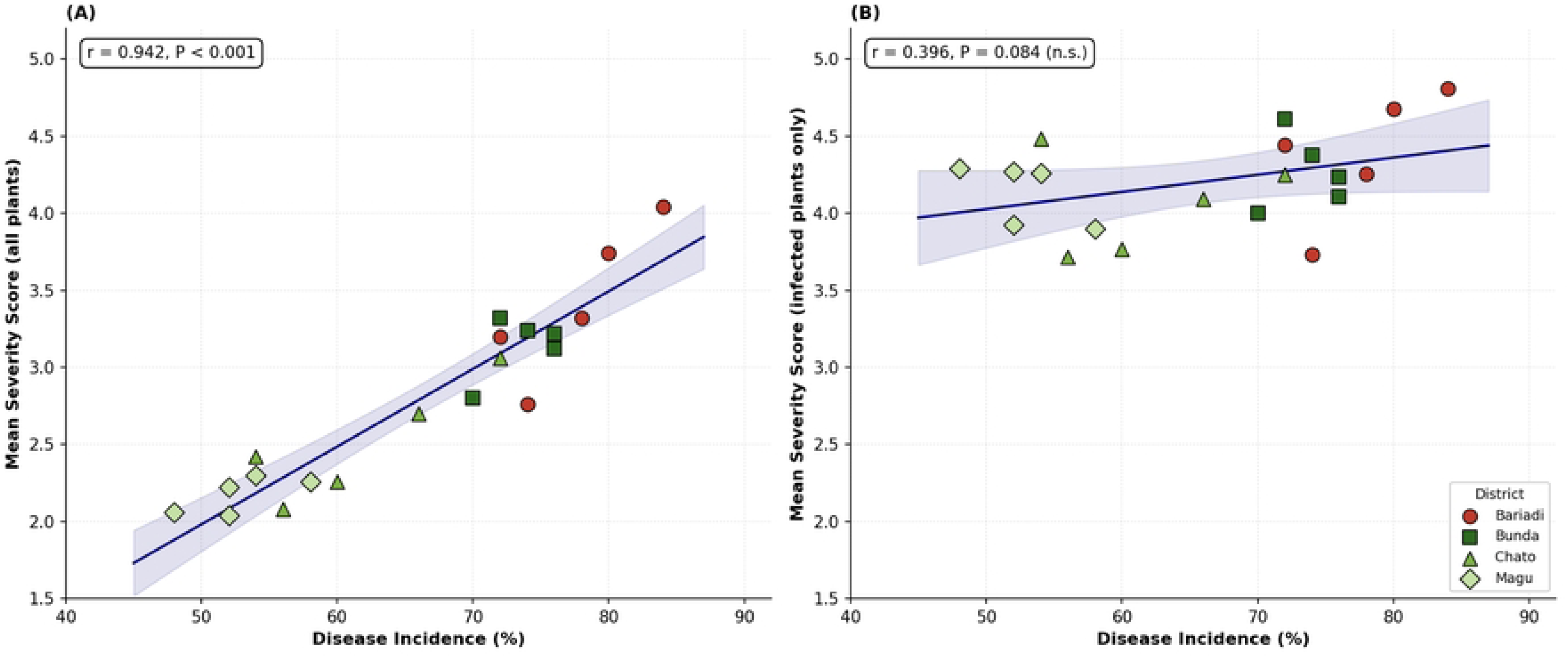
Relationship between disease incidence and severity of Alternaria leaf spot in cotton. (**A**) Correlation between disease incidence and mean severity score calculated using all assessed plants. (B) Correlation between disease incidence and mean severity score calculated using only infected plants, excluding healthy plants (severity class 0). Lines show fitted linear regressions with 95% confidence bands. n.s. = not significant.

### 3.6 Morphological identification of Alternaria isolates

A total of 40 fungal isolates were recovered from symptomatic cotton leaves collected across all 20 surveyed fields. All isolates exhibited colony and conidial characteristics consistent with the *genus Alternaria Nees*. Seven-day-old cultures on PDA displayed grayish to dark olive-gray colonies with velvety to woolly textures, irregular margins, and dense aerial mycelia; colony reverses were dark brown to blackish. Microscopic examination revealed dark brown, muriform, obclavate-to-ellipsoidal conidia with both transverse and longitudinal septa borne on simple or branched conidiophores, confirming genus-level identification.

Considerable morphological variation in conidia was observed among isolates. Conidial length ranged from 18.2 to 68.4 μm, width from 9.0 to 22.0 μm, transverse septa from 2 to 8, longitudinal septa from 0 to 3, and beak length from 5.0 to 95.0 μm. Despite this variation, all isolates retained the diagnostic cultural and microscopic features characteristic of the genus Alternaria. Representative photographs are shown in Figures 7 and 8. Definitive species identification requires multi-locus molecular sequence analysis (MLSA), which was beyond the scope of this study.

**Figure 7.**
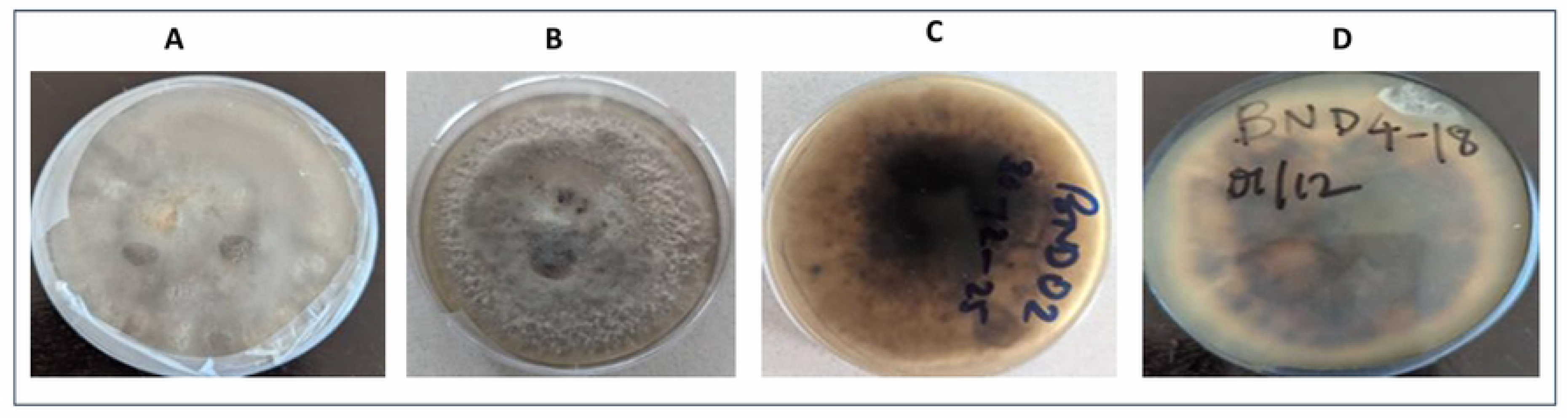
Colony morphology of representative Alternaria isolates recovered from cotton leaves on PDA after 7 days at 25°C. (A&B) Surface view showing greyish to dark olive-grey colonies. (C&D) Reverse view showing dark brown to blackish pigmentation

**Figure 8:**
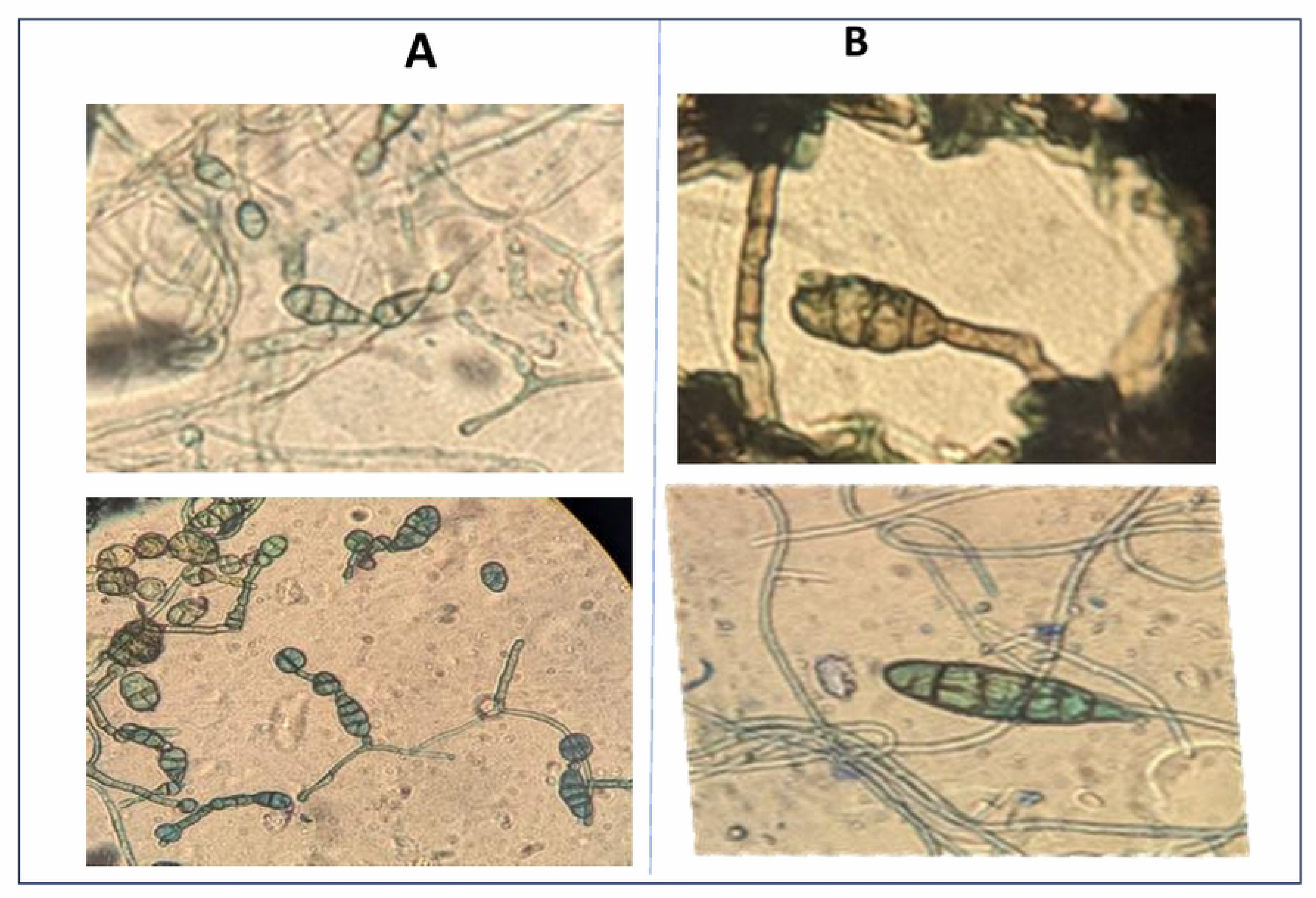
Microscopic characteristics of Alternaria conidia isolated from symptomatic cotton leaves collected from surveyed districts. (A) Shorter conidia in chains (B) Obclavate conidia with transverse and longitudinal septa and long filiform beak

## 4.0 DISCUSSION

The detection of Alternaria leaf spot in all 20 surveyed fields indicates that the disease is now widespread and well established across the main cotton-growing areas of the Lake Zone. However, this widespread presence at the field level does not imply uniform disease pressure: mean incidence (64.4%) and severity (DSI = 56.7%) varied markedly among districts. The correlation between incidence and severity weakened when the analysis was limited to infected plants, suggesting that the overall pattern was driven more by a higher proportion of infected plants than by increased severity among infected plants. Overall, these results indicate that while Alternaria leaf spot has become a significant, almost universal challenge to cotton farming in the Lake Zone, its impact varies geographically, influenced by differences in pathogen persistence, host presence, and environmental conditions conducive to disease development.

Documented yield losses of 30-40% in the Lake Zone under favourable conditions (Mung’ong’o, 2024; URT, 2023), coupled with 100% field prevalence, establish Alternaria leaf spot as a major production constraint that directly contributes to Tanzania’s documented productivity deficit of 1.34 tonnes/hectare versus the potential 3 tonnes/hectare (Ministry of Agriculture, 2023). Quantitative evidence from other cotton-producing regions corroborates these impact estimates. Under favourable conditions, Alternaria leaf spot causes yield reductions of 26.59% (Monga *et al*., 2013) and 38.23% (Bhattiprolu & Prasada Rao, 2009). Disease severity in comparable international surveys ranges from 2.16 to 24.12% in India (AICRP on Cotton, 2016-17) to up to 87.0% in New Mexico (Zhu *et al*., 2019). The substantially higher mean DSI in the Lake Zone (56.7%) than most international benchmarks, approaching the upper range reported in New Mexico’s high-incidence cotton regions, confirms that the Lake Zone represents a particularly high-risk agroecological environment for Alternaria leaf spot.

Widespread occurrence and epidemic development of Alternaria leaf spot have been reported across major cotton-growing regions worldwide, particularly in environments with favourable moisture and prolonged humidity. Studies by Monga *et al*. (2013), Lawrence *et al*. (2015), and Sampathkumar et al. (2024) consistently show that Alternaria epidemics intensify under conditions conducive to pathogen survival, sporulation, and secondary inoculum dispersal. The disease is considered an opportunistic foliar pathogen that frequently colonizes cotton plants during periods of physiological stress or favourable environmental conditions (Zhu *et al*., 2019; Wang *et al*., 2024). The consistent documentation of this pathogen across geographically diverse cotton-growing systems confirms that Alternaria leaf spot is a globally recurrent production constraint wherever environmental conditions align with the pathogen’s infection requirements, and provides a strong comparative basis for interpreting the epidemic levels observed in the Lake Zone.

The persistence and spread of Alternaria inoculum within the Lake Zone are further reinforced by agronomic practices and crop management systems commonly used by smallholder farmers in the region. Alternaria inoculum is well established within local cotton production systems and may persist between cropping seasons through infected crop residues and airborne spores. Continuous cotton cultivation with limited crop rotation likely facilitates the accumulation and survival of Alternaria inoculum in infected crop residues and volunteer cotton plants. Retaining infected cotton stover between growing seasons may provide favourable substrates for pathogen survival and subsequent epidemic initiation during the following cropping cycle, as previously reported in cotton production systems affected by Alternaria leaf spot (Zhu *et al*., 2019; Hu *et al*., 2016). Additionally, seedborne inoculum may contribute to long-distance dissemination of Alternaria spp., particularly in smallholder production systems where farmer-saved seed is widely used (Mohamed *et al*., 2019). Moreover, potassium deficiency, which may occur in nutrient-limited soils of the Lake Zone, could accelerate leaf senescence and increase susceptibility to Alternaria infection (Zhao *et al*., 2013; Naraghi *et al*., 2024).

The district-level rainfall data collected during the survey period suggest a plausible, though correlational, explanation for the observed spatial gradient in disease severity. Bariadi and Bunda, the districts with the highest incidence and severity, consistently recorded higher rainfall than Chato and especially Magu across most months of the 2024-2025 growing season, including the vegetative-to-boll development phase when disease assessments were conducted. This pattern aligns with the well-established understanding that leaf wetness, rainfall frequency, and humidity facilitate the germination, sporulation, and secondary spread of Alternaria conidia (Bhat *et al*., 2013; Kumar *et al*., 2018). However, this study did not statistically analyze rainfall, temperature, or humidity as predictors of disease incidence or severity; thus, the climatic association is descriptive rather than confirmatory. Other factors that can influence inter-district disease levels, such as cropping history, cultivar susceptibility, local crop management, and background inoculum load, were not measured and should be explored in future, more focused epidemiological research.

The morphological variation among the 40 isolates, with overlapping conidial dimensions, suggests that mixed infections may occur within the Lake Zone cotton-growing system, consistent with reports from other African and Asian cotton-producing regions (Sampathkumar *et al*., 2024; Lin *et al*., 2024). The morphological data support genus-level identification of all isolates as *Alternaria spp*., but definitive species-level attribution and pathogenicity confirmation will require multi-locus molecular sequence analysis (MLSA) of housekeeping genes such as glyceraldehyde-3-phosphate dehydrogenase (GAPDH) and controlled inoculation studies, as successfully applied in recent Alternaria species delineation studies (Woudenberg *et al*., 2013).

These findings provide important guidance for managing Alternaria leaf spot in the cotton-growing systems of Tanzania’s Lake Zone, particularly because disease development appears strongly influenced by environmental conditions, infected crop residues, and continuous cotton cultivation. Cultural management practices such as field sanitation, removal and destruction of infected cotton residues, crop rotation, and use of clean seed may help reduce inoculum buildup and subsequent epidemic development. Similar residue management practices have been successfully used to minimize the survival and spread of foliar fungal pathogens in major cotton-growing regions worldwide (Venkatesh *et al*., 2015). The observed distribution of the disease across districts and villages may further assist farmers and extension officers in identifying high-risk production areas that require intensified disease monitoring and early management interventions. Integrated disease management approaches that combine cultural practices, continuous disease surveillance, and climate-informed management strategies will be essential for reducing Alternaria leaf spot pressure and improving cotton productivity within Tanzania’s Lake Zone.

## 5.0 STRENGTHS AND LIMITATIONS OF THE STUDY

This study provides the first systematic epidemiological baseline on Alternaria leaf spot incidence and severity across four major cotton-growing districts in the Lake Zone of Tanzania, filling a critical knowledge gap. The use of a standardized diagonal sampling protocol, consistent disease assessment scales, and descriptive integration of district-level seasonal climatic data from the Tanzania Meteorological Authority strengthens the contextual interpretation of the findings.

Several limitations must be acknowledged. The cross-sectional design, involving a single growing season (2024-2025), limits the ability to capture interannual variability in disease dynamics. The seasonal climatic data presented in this study are descriptive and serve to characterize the environmental conditions prevailing during the survey period. Future research should employ time-series analysis across multiple growing seasons to quantify the predictive value of climate variables on Alternaria epidemiology. Molecular characterization of isolates using multi-locus sequence analysis and pathogenicity testing under controlled conditions to confirm the causal agent and assess potential differences in virulence were outside the scope of the present baseline survey.

## 6.0 CONCLUSION

Alternaria leaf spot (ALS) was found to be widespread across all surveyed cotton fields in the Lake Zone of Tanzania, with an overall mean incidence of 64.4% and a mean disease severity index (DSI) of 56.7%. Incidence and severity were significantly higher in Bariadi and Bunda than in Chato and Magu districts. All recovered isolates were morphologically consistent with the genus Alternaria.

This study provides the first systematic epidemiological baseline for Alternaria leaf spot across the major cotton-growing districts of the Lake Zone, Tanzania. Disease incidence and severity varied significantly among districts, highlighting the critical need for tailored disease management strategies. This study recommends continued surveillance and integrated disease management to safeguard cotton productivity among smallholder farmers in the region.

## ACKNOWLEDGEMENT

The authors gratefully acknowledge Sokoine University of Agriculture (SUA) for providing academic guidance and research support. Sincere thanks are extended to the Tanzania Agricultural Research Institute (TARI) for financial support and logistical facilitation. Appreciation is also expressed to the Tanzania Meteorological Authority (TMA) for providing climatic data, and to field technicians, District Agricultural, Livestock, and Fishery Officers (DALFOs), and cotton farmers of the Lake Zone of Tanzania, whose participation and cooperation made this research possible.

